# Data-driven hypothesis weighting increases detection power in multiple testing

**DOI:** 10.1101/034330

**Authors:** Nikolaos Ignatiadis, Bernd Klaus, Judith Zaugg, Wolfgang Huber

## Abstract

Hypothesis weighting is a powerful approach for improving the power of data analyses that employ multiple testing. However, in general it is not evident how to choose the weights in a data-dependent manner. We describe independent hypothesis weighting (IHW), a method for making use of informative covariates that are independent of the test statistic under the null, but informative of each test’s power or prior probability of the null hypothesis. Covariates can be continuous or categorical and need not fulfill any particular assumptions. The method increases statistical power in applications while controlling the false discovery rate (FDR) and produces additional insight by revealing the covariate-weight relationship. Independent hypothesis weighting is a practical approach to discovery of associations in large datasets.

## Introduction

Multiple testing is an important part of many high-throughput data analysis workflows. A common objective is control of the FDR, i. e., the expected fraction of false positives among all positives. Algorithms exist that achieve this objective by working solely off the list of p-values from the hypothesis tests [1–5]. However, such an approach tends to be suboptimal when the individual tests differ in their statistical properties, such as sample size, true effect size, signal-to-noise ratio, or prior probability of being false.

For example, in RNA-seq differential gene expression analysis, each hypothesis is associated with a different gene, and because of differences in the number of reads mapped per gene they may greatly differ in their signal-to noise ratio. In genome-wise association studies (GWAS), associations are sought between genetic polymorphisms and phenotypic traits; however, the power to detect an association is lower for rarer polymorphisms (all else being equal). In GWAS of gene expression phenotypes (eQTL), cis-effects are a priori more likely than associations between a gene product and a distant polymorphism.

To take into account the different statistical properties of the tests, one can associate each test with a weight, a non-negative number as a measure of its priority. The weights fulfill a budget criterion, commonly that they average to one. Hypotheses with higher weights get prioritized [6]. The procedure of Benjamini and Hochberg (BH) [1] can be modified to allow weighting simply by replacing the original p-values *p*_*i*_ with their weighted versions *p*_*i*_/*w*_*i*_ (where *w*_*i*_ is the weight of hypothesis *i*) [7]. However, FDR control of this approach is guaranteed only if the weights are pre-specified and thus independent of the data. In practice, the optimal choice of weights is rarely known, and a data-driven method would be desirable [8–12]. However, a method that is generally applicable, shows good performance and ensures type-I error control has been lacking.

## Results

Independent hypothesis weighting (IHW) is a multiple testing procedure that applies the weighted BH method [7] using weights derived from the data. The input to IHW is a two-column table of p-values and covariates. The covariate can be any continuous-valued or categorical variable that is thought to be informative on the statistical properties of the hypothesis tests, while it is independent of the p-value under the null hypothesis [10]. The latter property can be verified either mathematically [10] or empirically [13]. Simple diagnostic plots of the data can help assess these assumptions (Figure 1).

**Figure 1.**
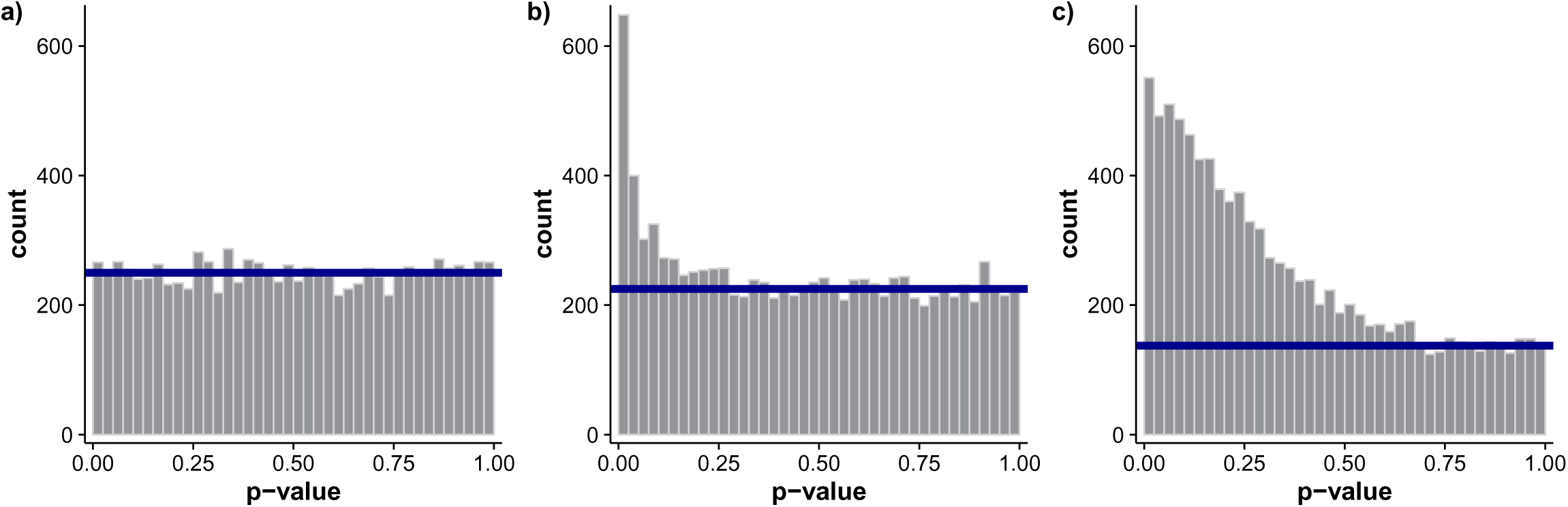
Stratified histograms as diagnostic plots. Histograms of the p-values after splitting the hypotheses by levels of the covariate help assess whether IHW is applicable. An approximately uniform distribution for larger values (e. g., *p* > 0.5) indicates that the p-values are well-calibrated irrespective of the covariate. The relative size of the peak to the left (compared to the uniform mixture component) and/or its shape should be different for different levels of the covariate. Here we show 3 exemplary histograms which illustrate these properties: **a**) Uniform p-values (no signal). **b**) Large proportion of nulls, small fraction of alternatives with strong signal. **c**) Smaller proportion of nulls, but a relatively weak signal for the alternatives.

We consider multiple testing as a resource allocation problem [7]: given a “budget” of acceptable FDR, how can it be distributed among the hypotheses in such a way as to obtain the best possible power overall? The first idea is to use information in available covariates. For instance, in differential expression analysis, a suitable covariate is the sum of read counts for the gene across all samples [10,13,14]; in expression-QTL or ChIP-QTL analysis, eligible covariates are the distance between the genetic variant and the genomic location of the phenotype, or measures of their comembership in a topologically associated domain [15]. Such covariates exist in many multiple testing applications. Therefore we assume that it is beneficial to assign hypothesis weights as a function of the covariate. Hypotheses with the same value of the covariate will be assigned the same weight. Furthermore, in our implementation we assume that the function can be approximated by a step-wise constant function, and note that no further assumptions (e. g., monotonicity) are needed.

The second idea is that the number of rejections of the weighted BH procedure with given weights is an empirical indicator of the method’s power. Therefore, a good choice of the covariate-to-weight mapping should lead to a high number of rejections.

Our first, naive implementation of IHW is easy to explain. Our algorithm divides the hypothesis tests into *G* different groups based on the covariate. Then, we associate each group with a weight, so that all hypotheses within one group *g* are assigned the same weight *w*_*g*_. For each possible weight vector w = (*w*_1_,…, *w*_*G*_) we apply the weighted BH procedure at level *α* and calculate the total number of rejections. We choose the weight vector **w^*^** that maximizes the number of rejections and report the corresponding result.

In many applications, this naive approach is already satisfactory, but it has two shortcomings: First, the underlying optimization problem is difficult and does not easily scale to problems with millions of hypothesis tests. Second, in certain situations, described below, this algorithm leads to loss of type I error control. These problems are overcome with three extensions of the naive algorithm:

E1. We find the optimal weight vector under a convex relaxation of the above optimization task, which in statistical terms corresponds to replacing the empirical cumulative distribution functions (ECDF) of the p-values with the Grenander estimators (least concave majorant of the ECDF). The resulting problem is convex and can be efficiently solved even for large numbers of hypotheses.

E2. We randomly split the hypotheses into k folds. For each fold, we apply convex IHW to the other *k* – 1 folds and assign the learned weights to the remaining fold. Thus the weight assigned to a given hypothesis does not directly depend on its p-value, but only on its covariate.

E3. The performance of the algorithm can be further improved by ensuring that the weights learned with *k* – 1 folds generalize to the held-out fold. Therefore, we introduce a regularization parameter *λ* ≥ 0, and the optimization is done over a constrained subset of the weights. For an ordered covariate, we require that 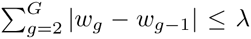, i.e., weights of successive groups should not be too different. For an unordered covariate, we use instead the constraint 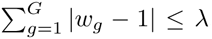, i.e., deviations from 1 are penalized. In the limit case *λ* = 0, all weights are the same, so IHW with *λ* = 0 is just the BH method. IHW with *λ* → ∞ is the unconstrained version. Choice of *λ* is a model selection problem, so within each split in E2 we apply a second nested layer of cross-validation. E3 is optional; whether or not to apply it will depend on the data. It will be most beneficial if the number of hypotheses per group is relatively small.

A complete description of the algorithm, including an efficient computational implementation of the optimization task, is provided in Supplemental Section 3. Supplemental Section 4 describes its theoretical justification.

### IHW increases empirical detection power compared to the BH procedure

We illustrate this claim on three exemplary applications. The first one, by Bottomly *et al*. [16,17], is a RNA-seq dataset used to detect differential gene expression between mouse strains. p-values were calculated using DESeq2 [13]. Here we used the mean of normalized counts for each gene, across samples, as the informative covariate. We saw an increased number of rejections compared to BH (Figure 2a). In addition, we observed that the learned weight function prioritized genes with higher mean normalized counts (Figure 2b).

**Figure 2.**
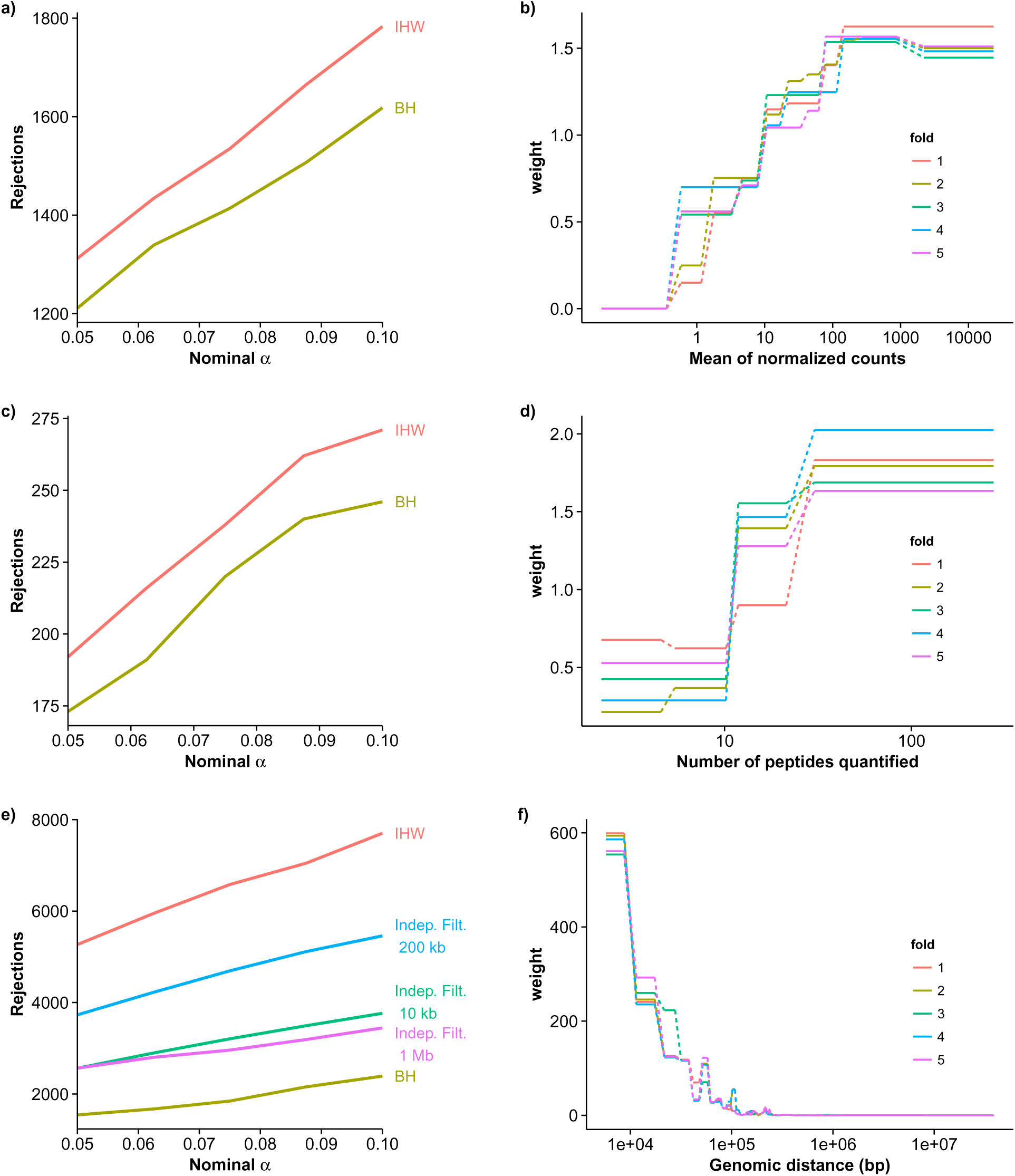
Comparison between IHW and BH. **a**) Benchmark based on Bottomly dataset [16], analyzed with DESeq2. IHW with the normalized mean of the counts for each gene as a covariate increases rejections compared to BH at different nominal significance levels. **b**) Learned weights at *α* = 0.1 for the Bottomly example. Genes with low counts get assigned low weight. Note that for each value of the covariate, five different weights have been learned because of the splitting scheme (E2). The trend is consistent. **c**) Rejections of IHW and BH for the SILAC dataset [18], plotted against the nominal significance level *α*. As a covariate we use the total number of peptides detected for each protein by the proteomics experiment. **d**) Weights learned by IHW at *α* = 0. 1 as a function of the total number of peptides quantified for each protein. **e**) Rejections of IHW, independent filtering with different thresholds and BH for Chromosome 21 of the hQTL dataset [15], plotted against the nominal significance level *α*. As a covariate we use the linear genomic distance between SNPs and ChIP-seq signals. IHW drastically increases the number of rejections. **f**) Weights learned by IHW at *α* = 0.1 as a function of genomic distance.

Second, we used a quantitative mass-spectrometry (SILAC) experiment in which yeast cells treated with rapamycin were compared to yeast treated with DMSO (2 × 6 biological replicates) [18]. Differential protein abundance of 2666 proteins was evaluated using Welch’s *t*-test [18]. As a covariate we used the total number of peptides which were quantified across all samples for each protein. IHW again showed increased power compared to BH (Figure 2c), and proteins whose peptides were quantified more often were assigned a higher weight, as expected (Figure 2d).

In a third example, we searched for associations between SNPs and histone modification marks (H3K27ac) [15] on human Chromosome 21. This yielded 180 million tests. As a covariate we used the linear genomic distance between the SNP and the ChIP-seq signal. As shown in Figure 2e, the power increase compared to BH and to independent filtering was dramatic. IHW automatically assigned most weight to small distances (Figure 2f). Thus IHW acted similarly to the common practice in eQTL-analysis of searching for associations only within a certain distance – but with the advantage that no arbitrary choice of distance threshold is needed, and that the weights are more nuanced than a hard distance threshold. IHW does not exclude SNP-phenotype pairs far away, and these can still be detected as long as they have a sufficiently small p-value.

### The extensions to naive IHW are needed to ensure type I error control

Naive IHW, as well as previous approaches to data-driven hypothesis weighting or filtering, do not maintain FDR control in situations where all hypotheses are true (Figure 3a) or where there is insufficient power to detect the false hypotheses (Figure S1a). In addition, the local fdr methods (Clfdr and FDRreg) often show strong deviations from the target FDR in a direction (conservative or anti-conservative) that is not apparent a priori (Figure 3b,c,e). Thus, among all methods benchmarked across these scenarios, only BH, IHW and LSL-GBH generally control the FDR. The results of our method comparisons are summarised in Table 1, and the simulations are described in Supplemental Section 7.

**Figure 3.**
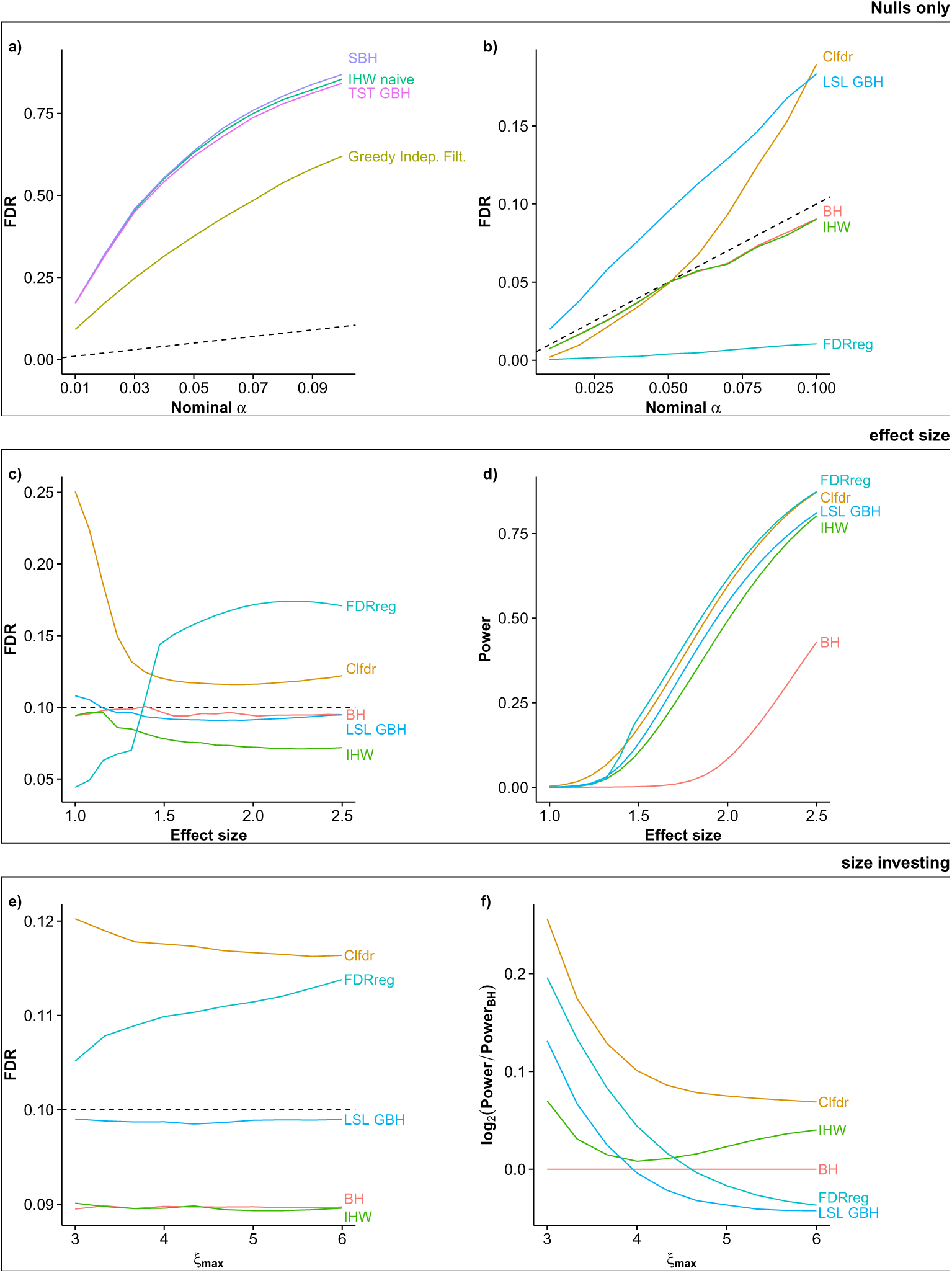
Assessment of methods’ performance in three different scenarios. Panels **a, b**) benchmark type I error control if all null hypotheses are true. Shown is the true FDR against the nominal significance level *α*; the dashed lines indicates the identity function. Brief descriptions of each method shown are in Table 1. **a**) Greedy independent filtering, SBH, TST GBH and naive IHW lose type I error control and make too many rejections. **b**) BH, FDRreg, and IHW control *α*, but for example note that FDRreg is too conservative in this case. LSL-GBH and Clfdr are slighly anticonservative, but a lot less than the methods in panel **a**). Panels **c, d**) look at implications of different effect sizes. The two-sample *t*-test was applied to Normal samples (*n* = 2 × 5, *σ*=1) with either the same mean (true nulls) or means differing by the effect size indicated on the *x*-axis (true alternatives). The fraction of true alternatives was 0.05. The pooled variance was used as the covariate. Rejections were made at the nominal level *α* = 0.1 (dotted line). **c**) The *y*-axis shows the actual FDR. Clfdr and FDRreg are strongly biased, and this bias depends on the true effect size. **d**) Power analysis. All methods show improvement over BH. Panels **e, f**) highlight size-investing. Data were simulated as in [8]: p-values were computed from the one-sided z-test, with *Z*_*i*_ ~ ***N***(0,1) under the null and *Z*_*i*_ ~ ***N***(*ξ*_*i*_, 1) under the alternative, with the covariate *ξ*_*i*_ ~ *U*[1, *ξ*_*max*_]. The fraction of alternatives was 0.1. **e**) The *y*-axis shows the actual FDR. In this case, Clfdr and FDRreg are slightly anticonservative. **f**) The logarithm of the power increase compared to BH is shown as a function of *ξ*_*max*_. Large values of *ξ*_*max*_ necessitate a size-investing strategy. Methods which cannot apply such a strategy, such as LSL-GBH and FDRreg, have power even lower than BH.

**Table 1.** Short description of the different methods benchmarked and summary of the results of Figure 3 and Figure S1.

| Method | Short description | Type I error control |  | Gain in power |  | Comment |
| --- | --- | --- | --- | --- | --- | --- |
| | | $\pi_0=1$ | t-test | t-test<br>(vs BH) | size<br>investing | |
| BH | Method of Benjamini and Hochberg [1] to control false-discovery rate (FDR) for multiple exchangeable hypotheses. | ✓ | ✓ | — | — |  |
| IHW | Independent hypothesis weighting, as proposed here. | ✓ | ✓ | ✓ | ✓ |  |
| Naive IHW | Naive independent hypothesis weighting, as proposed here. | ✗ | ✗ | ✓ | ✓ |  |
| Greedy independent filtering | The independent filtering procedure [10] modified to use a data-driven filter threshold which maximizes the number of rejections. | ✗ | ✗ | ✓ | ✗ | The covariate-weights function is a binary step, monotonic. |
| SBH | Stratified Benjamini-Hochberg [20]: Apply the BH procedure at level $\alpha$ within each stratum, then combine the rejections across the strata. | ✗ | ✗ | ✓ | ✗ | |
| TST-GBH | The Group BH procedure [11]: An adaptive weighted BH procedure applied with weights proportional to $\frac{\pi_1}{\pi_0}$ within each group. $\pi_0$ is estimated using the TST estimator [31]. | ✗ | ✗ | ✓ | ✗ | |
| LSL-GBH | The Group BH procedure [11], where $\pi_0$ is estimated using the LSL estimator [2]. | ✓ | ✓ | ✓ | ✗ | |
| Clfdr | In the CLfdr procedure [22], the local fdr is estimated separately within each group and the estimates are pooled together. For the fdr estimation here we use the modified Grenander estimator [5]. | ✓ | ✗ | ✓ | ✓ |  |
| FDRreg | The FDRregression method [25] estimates the local fdr by assuming all hypotheses have the same alternative density and $\pi_0$ varies smoothly as a function of the covariate. | ✓ | ✗ | ✓ | ✗ | Requires $z$-scores. |

### IHW can apply a size investing strategy

The term *size investing* refers to the notion that most of the type I error budget (’size’) of a multiple testing decision should be invested where it might make most of a difference: i. e., not into tests that are likely not to be rejected anyway, however high the weight they are given; nor into tests with very high power, so that rejections will be made anyway, even with a low weight; but rather, in the middle ground, where some additional weight might make the difference between rejection or not [19]. Size investing is provided by IHW but not by LSL-GBH and FDRreg, and as shown in Figure 3f, the latter might even lose power compared to the BH method because of that. A reason is that in the latter, weight differences are driven only by differences in *π*_1_ and not by the alternative distribution. This is further elaborated in the supplement. Also note that independent filtering [10] and stratified BH [20] cannot capture such a size-investing dynamic (Figure S1d).

### Relationship between IHW and local true discovery rates

Recall the motivation for IHW as a resource allocation problem. This was triggered by the observation that p-values lose information [21], in the sense that they do not capture important statistical properties of each test. Indeed, a better representation of these properties is provided by the concept of the *local true discovery rate* (tdr) [4]. The tdr of the *i*^th^ hypothesis is [4]

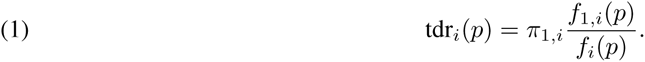

A schematic explanation is given in Figure 4a (see also Figure S2). *f*_*i*_ is the density of the distribution of the p-value *p*. It is a mixture of two conditional distributions, *f*_*i*_ = *π*_0, *i*_*f*_0_ + *π*_1, *i*_*f*_1, *i*_, where the densities *f*_0_ and *f*_1, *i*_ are conditional on the null or the alternative being true, respectively, and *π*_0, *i*_ and *π*_1, *i*_ (which sum up to 1) are the corresponding prior probabilities. The null distribution of a properly calibrated test is uniform, therefore we can set *f*_0_(*p*) = 1 irrespective of *p* and *i*. In Figure 4b–d three hypotheses are shown with different tdr curves corresponding to different power profiles.

**Figure 4.**
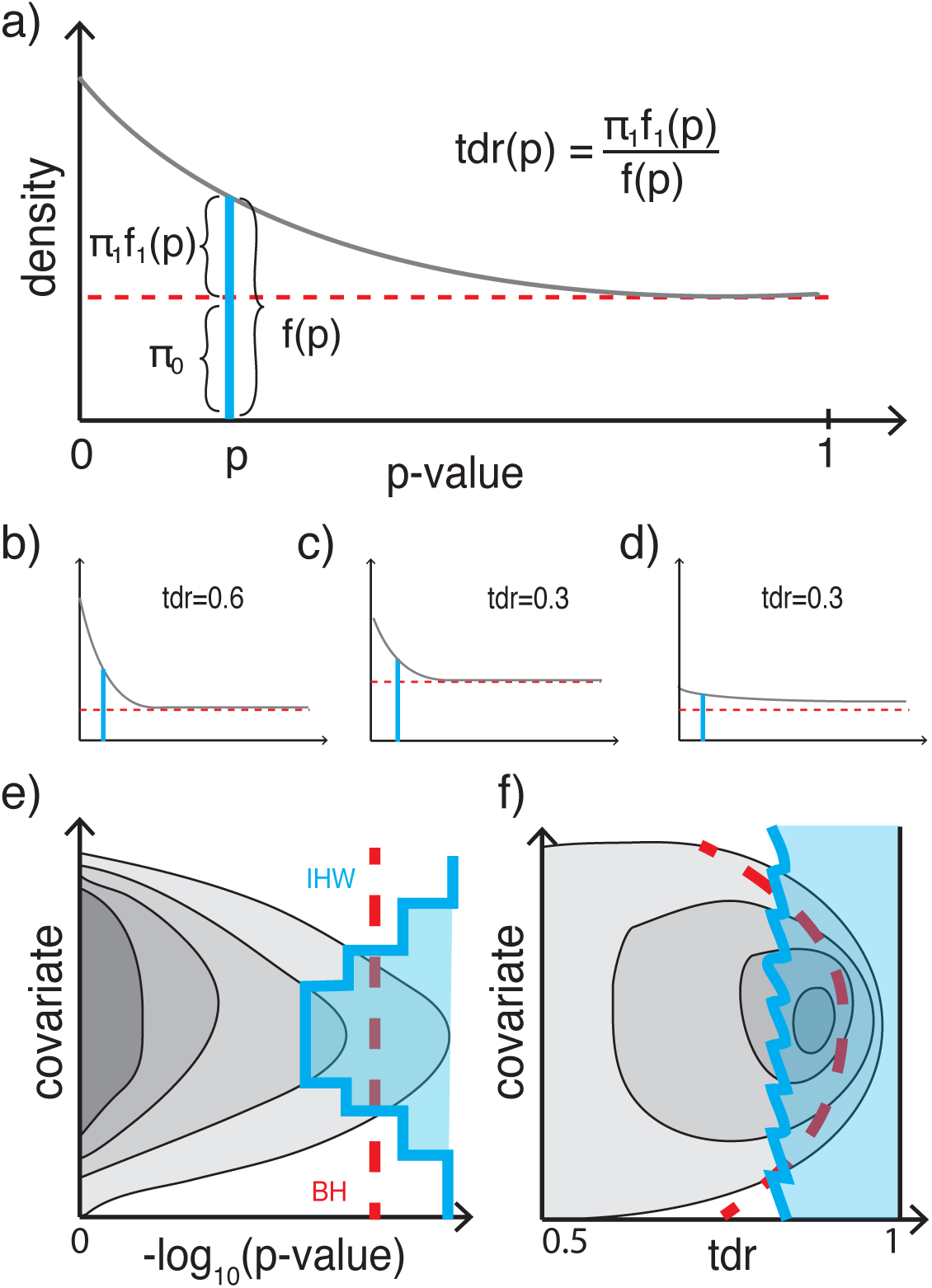
True discovery rate and informative covariates. **a**) Schematic representation of the mixture density *f*_*i*_, which is composed of the alternative density *f*_1, *i*_ weighted by its prior probability *π*_1, *i*_ and the uniform null distribution weighted by *π*_0, *i*_. **b-d**) The true discovery rate (tdr) of individual tests can vary for different reasons. In **b**), the test has high power, and the prior probability of the null is well below 1. In **c**), the test has equally high power, but the the prior probability of the null is higher, leading to a reduced tdr. In **d**), the prior probabilities are like in **b**), but the test has little power, again leading to a reduced tdr. **e**) If an informative covariate is associated with each test, the distribution of the p-values from multiple tests is different for different values of the covariate. The contours schematically represent the joint two-dimensional density of p-values and covariate. The rejection boundaries of the BH procedure do not take into account covariates and only depend on the p-values (dashed red line). In contrast, the decision boundary of IHW is a step function; each step corresponds to one group, i. e., to one weight. **f**) By virtue of Equation (1), the density of the tdr also depends on the covariate. The decision boundary of the BH procedure is shown by the dashed red line; it leads to a suboptimal set of rejections, in this example with lower than optimal tdr for small and large values of the covariate, and higher than optimal tdr for intermediate values. In contrast, the decision boundary of IHW approximates a line of constant tdr, implying a near-optimal use of the type I error budget to make as many rejections as possible. An important feature of the IHW method is that it does not require explicit estimation of the two-dimensional densities, but instead works directly on the p-values and the covariate.

It can now be shown that to maximize power at a given FDR, one should reject the hypotheses with the highest tdr_*i*_(*p*_*i*_) [22,23]. In other words, if we knew the functions tdr_*i*_ and could use tdr_*i*_(*p*_*i*_) as our test statistics, then without any further effort we would have a method for FDR control with optimal power.

Similarly to the central idea of IHW, one can now assume that the many different, unknown univariate functions tdr_*i*_(*p*), one for each hypothesis *i*, can be usefully approximated by a single bivariate function tdr(*p*, *x*), where *x* is the covariate. Then contour lines of the joint density of tdr and *x* are as illustrated in Figure 4f. We can see how in such a scenario the rejection region of the BH method tends to be suboptimal. As it is defined solely in terms of p-values (Figure 4e), it differs from the optimal region, whose boundary is a vertical line of constant tdr (Figure 4f).

However, in practice, we do not usually know the quantities in Equation (1) or the bivariate function tdr(*p*, *x*) and have to estimate them [24]. Unfortunately, this estimation problem is difficult, and even with the use of additional approximations, such as splines [25] or piecewise constant functions [26], there does not seem to be a practical implementation.

On the other hand, with IHW we circumvent explicit estimation of the bivariate tdr function and instead derive a powerful testing procedure by assigning data-driven hypothesis weights. In addition, the IHW method readily extends to other weighted multiple testing testing procedures [7]. In Supplemental Section 5, we describe IHW-Bonferroni, a new powerful method for control of the familywise error rate (FWER). In contrast, local tdr methods are specific to the FDR.

## Discussion

We have introduced a weighted multiple testing method that learns the weights from the data. Its appeal lies in its generic applicability. It does not require assumptions about the relationship between the covariate and the power of the individual tests, such as monotonicity, which is necessary for independent filtering. It can apply size investing strategies, since it does not assume that the alternative distributions are the same across the different hypotheses. It is computationally robust and scales to millions of hypotheses.

The idea of using informative covariates for hypothesis weighting or for shaping optimal rejection boundaries is not new (Table 1; [26–29]). In this work, we provide a general and practical approach. Most importantly, we show how to overcome two major limitations of previous approaches: type I error control and stability. To retain FDR control, we introduced a strategy reminiscent of Tibshirani’s and Efron’s prevalidation procedure [30], in which we randomly split the hypotheses into folds and then assign weights to a given hypothesis based on training on the other folds. To ensure convexity of the optimisation problem and computational scalability, we replaced the empirical distribution function by the Grenander estimator. We have observed that this extension alone already reduces the overoptimism of naive IHW to a large extent, even without the splitting scheme (E2).

Various approaches to increasing power compared to BH have focused on estimating the value of *π*_0_, the fraction of true nulls among all hypotheses, in cases where it is less than 1, as the BH method uses the conservative upper limit [2,31]. In practice, this tends to have limited impact, since in the most interesting situations the number of true discoveries is small compared to all tests, so that *π*_0_ is close to 1 [32] and no substantial power increase is gained. On the other hand, for IHW this situation could be changed, since often the groups that get assigned a high weight also have an increased *π*_1_.

In our method we have explicitly avoided estimating the densities in Equation (1). Nevertheless, the local tdr is an interesting quantity in its own right, since it provides a posterior probability for each individual hypothesis. Our weighted p-values do not provide this information. Thus, development of stable estimation procedures for the local tdr that incorporate informative covariates is needed and would be complementary to our work [24–26,33].

### Software availability

The IHW package, which implements the proposed multiple testing procedure, is available on Bioconductor [34]. It comes with detailed documentation and a vignette that showcases the application of IHW to a real dataset. The vignette also provides theoretical guidance on the choice of informative covariates and suggests diagnostic plots, so that users can determine if their covariate satisfies the required conditions.

Executable documents (Rmarkdown) reproducing all analyses shown here can be downloaded at https://github.com/nignatiadis/IHWpaper.

## Acknowledgements

We thank Bernd Fischer, Michael I. Love, Mikhail Savitski and Oliver Stegle for insightful discussions and comments on the manuscript, the COIN-OR project for the open-source *SYMPHONY* software, and Vladislav Kim for interfacing it to R through the *lpsymphony* package. We acknowledge support from the European Commission through the Horizon 2020 project SOUND.

## REFERENCES

[1] Benjamini, Y. & Hochberg, Y. Controlling the false discovery rate: a practical and powerful approach to multiple testing. Journal of the Royal Statistical Society. Series B (Methodological) 289–300 (1995).

[2] Benjamini, Y. & Hochberg, Y. On the adaptive control of the false discovery rate in multiple testing with independent statistics. Journal of Educational and Behavioral Statistics 25, 60–83 (2000).

[3] Storey, J. D., Taylor, J. E. & Siegmund, D. Strong control, conservative point estimation and simultaneous conservative consistency of false discovery rates: a unified approach. Journal of the Royal Statistical Society: Series B (Statistical Methodology) 66, 187–205 (2004).

[4] Efron, B. Large-scale inference: empirical Bayes methods for estimation, testing, and prediction (Cambridge University Press, 2010).

[5] Strimmer, K. A unified approach to false discovery rate estimation. BMC Bioinformatics 9, 303 (2008).

[6] Holm, S. A simple sequentially rejective multiple test procedure. Scandinavian Journal of Statistics 65–70 (1979).

[7] Genovese, C. R., Roeder, K. & Wasserman, L. False discovery control with p-value weighting. Biometrika 93, 509–524 (2006).

[8] Roeder, K., Devlin, B. & Wasserman, L. Improving power in genome-wide association studies: weights tip the scale. Genetic epidemiology 31, 741–747 (2007).

[9] Roquain, E. & Van De Wiel, M. Optimal weighting for false discovery rate control. Electronic Journal of Statistics 3, 678–711 (2009).

[10] Bourgon, R., Gentleman, R. & Huber, W. Independent filtering increases detection power for high-throughput experiments. Proceedings of the National Academy of Sciences 107, 9546–9551 (2010).

[11] Hu, J. X., Zhao, H. & Zhou, H. H. False discovery rate control with groups. Journal of the American Statistical Association 105 (2010).

[12] Dobriban, E., Fortney, K., Kim, S. K. & Owen, A. B. Optimal multiple testing under a gaussian prior on the effect sizes. Biometrika 102, 753–766 (2015).

[13] Love, M. I., Huber, W. & Anders, S. Moderated estimation of fold change and dispersion for RNA-Seq data with DESeq2. Genome Biology 15, 550 (2014).

[14] Robinson, M. D., McCarthy, D. J. & Smyth, G. K. edgeR: a Bioconductor package for differential expression analysis of digital gene expression data. Bioinformatics 26, 139–140 (2010).

[15] Grubert, F. et al. Genetic control of chromatin states in humans involves local and distal chromosomal interactions. Cell 162, 1051–1065 (2015).

[16] Bottomly, D. et al. Evaluating gene expression in C57BL/6J and DBA/2J mouse striatum using RNA-Seq and microarrays. PloS one 6, e17820 (2011).

[17] Frazee, A. C., Langmead, B. & Leek, J. T. Recount: a multi-experiment resource of analysis-ready rna-seq gene count datasets. BMC bioinformatics 12, 449 (2011).

[18] Dephoure, N. & Gygi, S. P. Hyperplexing: a method for higher-order multiplexed quantitative proteomics provides a map of the dynamic response to rapamycin in yeast. Science signaling 5, rs2–rs2 (2012).

[19] Peña, E. A., Habiger, J. D. & Wu, W. Power-enhanced multiple decision functions controlling family-wise error and false discovery rates. The Annals of Statistics 39, 556–583 (2011).

[20] Yoo, Y. J., Bull, S. B., Paterson, A. D., Waggott, D. & Sun, L. Were genome-wide linkage studies a waste of time? exploiting candidate regions within genome-wide association studies. Genetic Epidemiology 34, 107–118 (2010).

[21] Sun, W. & Cai, T. T. Oracle and adaptive compound decision rules for false discovery rate control. Journal of the American Statistical Association 102, 901–912 (2007).

[22] Cai, T. T. & Sun, W. Simultaneous testing of grouped hypotheses: Finding needles in multiple haystacks. Journal of the American Statistical Association 104 (2009).

[23] Ochoa, A., Storey, J. D., Llins, M. & Singh, M. Beyond the E-value: Stratified statistics for protein domain prediction. PLoS Comput Biol 11, e1004509 (2015).

[24] Ploner, A., Calza, S., Gusnanto, A. & Pawitan, Y. Multidimensional local false discovery rate for microarray studies. Bioinformatics 22, 556–565 (2006).

[25] Scott, J. G., Kelly, R. C., Smith, M. A., Zhou, P. & Kass, R. E. False discovery rate regression: an application to neural synchrony detection in primary visual cortex. Journal of the American Statistical Association 110, 459–471 (2015).

[26] Ferkingstad, E., Frigessi, A., Rue, H., Thorleifsson, G. & Kong, A. Unsupervised empirical Bayesian multiple testing with external covariates. The Annals of Applied Statistics 714–735 (2008).

[27] Efron, B. & Zhang, N. R. False discovery rates and copy number variation. Biometrika 98, 251–271 (2011).

[28] Pickrell, J. K. Joint analysis of functional genomic data and genome-wide association studies of 18 human traits. The American Journal of Human Genetics 94, 559–573 (2014).

[29] Du, L. & Zhang, C. Single-index modulated multiple testing. The Annals of Statistics 42, 30–79 (2014).

[30] Tibshirani, R. J. & Efron, B. Pre-validation and inference in microarrays. Statistical Applications in Genetics and Molecular Biology 1 (2002).

[31] Benjamini, Y., Krieger, A. M. & Yekutieli, D. Adaptive linear step-up procedures that control the false discovery rate. Biometrika 93, 491–507 (2006).

[32] Benjamini, Y., Heller, R. & Yekutieli, D. Selective inference in complex research. Philosophical Transactions of the Royal Society A: Mathematical, Physical and Engineering Sciences 367, 4255–4271 (2009).

[33] Stephens, M. False discovery rates: A new deal. bioRxiv 038216 (2016).

[34] Huber, W. et al. Orchestrating high-throughput genomic analysis with bioconductor. Nature methods 12, 115–121 (2015).

